# Human immune response to primary cryptosporidiosis parallels murine infection models

**DOI:** 10.1101/2025.10.03.680261

**Authors:** Dana Van Fossen, Haroldo J. Rodriguez, Farha Naz, Carol A. Gilchrist, Justin J. Taylor, William A. Petri, Audrey C. Brown

## Abstract

*Cryptosporidium* is a protozoan parasite that causes cryptosporidiosis, an enteric infection associated with diarrhea, malnutrition, and impaired childhood development in low- and middle-income countries. Both humoral and cell-mediated immune responses have been implicated in protection, but the durability and quality of human immune responses in immunocompetent adults remain poorly defined. We investigated the development of immunity in two healthy US adults following primary cryptosporidiosis acquired during travel to Bangladesh. Longitudinal plasma samples were analyzed for antibody responses to *Cryptosporidium* antigens Cp17 and Cp23 and for circulating cytokine profiles. Circulating antibody peaked at three weeks post-infection but declined rapidly thereafter, approaching baseline within 16 weeks. In contrast, antibody avidity increased steadily over time, consistent with ongoing affinity maturation in germinal centers. While affinity maturation occurred, the composition memory B cells specific to *Cryptosporidium* antigens was skewed toward IgM+ cells across timepoints suggesting extrafollicular responses dominated and germinal center-derived, class-switched memory was limited. Cytokine profiling revealed an acute Th1-skewed response, with elevations in CXCL9, CXCL10, IL-27, IFNγ, IL-12, and IL-18 during early infection. These signatures mirrored protective pathways identified in murine models, underscoring the importance of type I immunity in parasite clearance. Together, these findings highlight that while antibody responses to *Cryptosporidium* are short-lived, avidity maturation persists, and Th1-driven cytokine responses dominate during acute infection. This work provides rare longitudinal data on immune responses in naïve adults following natural cryptosporidiosis and offers insight into mechanisms that may inform vaccine development and strategies to mitigate recurrent infection in vulnerable populations.

## Introduction

*Cryptosporidium* is a protozoan parasite that causes cryptosporidiosis, an enteric infection characterized by watery diarrhea and, in some cases, severe dehydration and malnutrition. The burden of disease is particularly high in low- and middle-income countries, where repeated infections in early childhood have been associated with growth faltering, impaired cognitive development, and increased mortality (1-5). Despite its global importance, our understanding of the immune responses that mediate protection and resolution of *Cryptosporidium* infection in humans remains incomplete.

Both cell-mediated and humoral immune mechanisms are implicated in defense against *Cryptosporidium*. Epidemiological and experimental studies indicate that T cell mediated immunity is essential for parasite clearance (6-8). This T cell mediated protection against cryptosporidiosis is largely driven by a type I (Th1) cytokine response, namely IFNγ, IL-12, and TNFa (6-8).

Available data suggest B cell responses and antibodies also contribute to limiting reinfection (6-8). Individuals who are deficient in B cell class-switch recombination, commonly due to mutations in the gene encoding CD40 ligand (CD40L), are susceptible to chronic infections with *Cryptosporidium* (9, 10). Fecal antibodies against the *Cryptosporidium* antigens Cp17 and Cp23 have been associated with delayed time to reinfection and a decline in the frequency of *Cryptosporidium*-associated diarrhea (11-14). However, the durability and functional quality of antibody responses following a natural infection in the absence of subsequent re-exposure are not well defined. Similarly, although cytokine production has been studied in animal models, human data on the kinetics of cytokine responses during acute infection and convalescence remain limited.

To address these gaps, we studied immune responses in two healthy adults with primary cryptosporidiosis in the absence of subsequent re-exposure. Longitudinal plasma and peripheral blood mononuclear cells (PBMCs) samples were collected to assess both 1) humoral immunity through antibody quantity and avidity over time and through immunoglobulin subclass phenotyping of memory B cells against Cp17 and Cp23 antigens and 2) cell-mediated immunity inferred though circulating cytokine levels. Our analyses revealed that antibody levels against *Cryptosporidium* Cp17 and Cp23 antigens peaked three weeks from infection but declined rapidly after disease resolution. The composition memory B cells specific to *Cryptosporidium* antigens was skewed toward IgM+ cells across timepoints. Plasma cytokine profiling demonstrated an acute IFNγ-dependent response associated with T-helper 1 cell and cytotoxic T cell activation, suggesting that cell-mediated immunity plays a central role in controlling infection. Understanding the relative contributions and limitations of these immune pathways may inform vaccine development and strategies to prevent recurrent infections in vulnerable populations.

## Methods

### Study approval and sample collection

This study was approved by the Institutional Review Board of the University of Virginia (IRB# HSR220013). Informed written consent was obtained from all participants in this study. Blood samples were drawn in EDTA Blood Collection Tubes (BD, Franklin Lakes, NJ). Plasma was obtained by sample centrifuging at 2000xG for 15 min, removing of the top liquid layer, and storing in aliquots at -70C. Then, the three times the volume of PBS plus 2% fetal bovine serum (FBS) as plasma removed was added to the blood collection tube and mixed by inversion. The diluted sample was layered over Lymphoprep (STEMCELL Technologies, Vancouver, BC), centrifuged at 450xg for 30 min, and the peripheral blood mononuclear cell (PBMC) band was collected. PBMCs were washed with PBS plus 2% FBS. The cell pellet was resuspended in RPMI plus 10% FBS followed by an equal volume of 2X cryopreservation media before freezing at -70C followed by long term storage in liquid nitrogen.

### Cryptosporidium antibody measurements

Enzyme-linked immunosorbent assay (ELISA) plates were coated with 5ug/mL Cp17 or Cp23 protein overnight at 4C. Plates were washed three times with PBS mixed with 0.05% Tween 20, blocked at 37C for 1 hr with ELISA blocking solution (Thermo Fisher Scientific, Waltham MA), and washed (x3) again. Plasma samples diluted 1:150 or 1:1500 were added in quadruplicate for 1 hr at 37C. Following a wash (x3), two wells of each sample were treated with 0.5M NH4SCN at room temperature for 15 min. An additional wash (x3) was performed, then HRP-conjugated anti-human IgM, IgA, or IgM was added for 1 hr at 37C. After a final wash (x3), TMB Turbo ELISA Substrate (Thermo Fisher Scientific, Waltham MA) was added for 7.5 min. 2N H2SO4 was used to stop the reaction before spectrophotometric measurement at 450nm was read and corrected with absorbance at 570 nm.

### Antigen tetramers and flow cytometry

Antigen tetramers were made for identification of antigen-specific B cells as previously described by Phelps, *et al*. (15). In brief, recombinant Cp17 and Cp23 proteins (Genscript, cat. no. For RQ-0000128619) were biotinylated with an EZ-Link Sulfo–NHS–LC–Biotinylation kit (Thermo Scientific, cat. no. 21435). Unreacted Sulfo-NHS-biotin was removed via size exclusion column before incubation with fluorophore-labeled streptavidin (SA-PE for Cp23, SA-APC for Cp17) (Agilent, cat. no. PJRS25; Cedarlane, cat. no. PJ27S). To exclude B cells binding SA, PE, and APC, control ‘decoy’ tetramers were generated in parallel with Cp17 and Cp23 tetramers. Recombinant Cp17 and Cp23 proteins contained an irrelevant HIS-tag protein at their N-terminus, so biotinylated His-tagged control protein was conjugated to an additional fluorophore-labeled streptavidin (SA-PE-DL594 or SA-APC-755) (Thermo Scientific, cat. no. 62266 and 62279) to provide a unique spectral fingerprint from the antigen tetramers.

For Cp17 and Cp23 B cell phenotyping, PBMCs from weeks 1, 3, 5, and 16 were thawed in warm RPMI supplemented with 10% fetal bovine serum (FBS) and immediately incubated in PBS with Live/Dead Blue (FVD eF506). Cells were then incubated for 10 minutes in FACS buffer (2% FBS, 0.5 mM EDTA) with human TruStain FcX Fc receptor block (Biolegend, cat. no. 422301), both Cp17 and Cp23 antigen tetramers, and decoy tetramers. All remaining surface markers, seen in **Table 1**, were then added and incubated for 30 minutes. Stained cells were washed before immediate acquisition by spectral flow cytometry (Cytek Aurora Borealis), and subsequent data was normalized by subsampling to an equal total live cell population across timepoints.

**Table 1.** Reagents used for flow cytometry analysis of PBMCs.

| **Marker** | **Fluorochrome** | **Company** | **Catalog** |
| --- | --- | --- | --- |
| FVD eF506 | BV510 | ThermoFisher | 65-0866-18 |
| CD3 | BV711 | BD | 563725 |
| CD14 | BV711 | BD | 563372 |
| CD16 | BV711 | BD | 563127 |
| CD20 | BV421 | BD | 740002 |
| CD19 | PE-Cy7 | BD | 560728 |
| CD21 | RY703 | BD | 771379 |
| CD11c | BUV395 | BD | 563787 |
| CD38 | BUV737 | BD | 612824 |
| CD138 | BUV496 | BD | 749874 |
| CD27 | VF563 | ThermoFisher | 365-0279-41 |
| IgA | BUV805 | BD | 772212 |
| IgG | BV605 | BD | 563246 |
| IgM | FITC | BD | 555782 |
| IgD | PerCP-Cy5.5 | BD | 561315 |
| CD71 | BV786 | BD | 563768 |
| Cp23 Tetramer | PE | N/A | N/A |
| PE HIS Decoy | PE-DL594 | N/A | N/A |
| Cp17 Tetramer | APC | N/A | N/A |
| APC HIS Decoy | APC-DL755 | N/A | N/A |

\*N/A indicates tetramer/fluorochrome combinations made in house

### Cytokine and biomarker quantification

C-reactive protein was measured in plasma diluted 1:200 using an ELISA kit according to the manufacturer’s instructions (ALPCO Diagnostics, Salem, NH). All other cytokine measurements were conducted using a multiplex Luminex bead-based assay (Luminex Corp., Austin, TX) according to the manufacturer’s protocol at the UVA Flow Cytometry Core Facility.

## Results

### Case description

In December 2024, a 30-year-old female (Subject A) and an 18-year-old female (Subject B), presented separately to outpatient clinics in Charlottesville, VA with watery diarrhea. Subjects A and B had recently traveled together to Bangladesh and returned to the USA one week prior to diarrheal symptom onset. Within Bangladesh, the patients visited rural areas of the Dinajpur and Tangail districts and urban communities within the capital, Dhaka.

PCR-based stool testing for gastrointestinal pathogens was positive for *Cryptosporidium spp*. in both patients. Subject A was additionally positive for enteropathogenic *Escherichia coli* (EPEC). Subject B was additionally positive for enterotoxigenic *Escherichia coli* (ETEC) and a Shiga-like toxin producing organism. Following exposure, the incubation periods for EPEC, ETEC, and Shiga-like toxin producing *E. coli* are 1-6, 1-3, and 3-8 days, respectively (16). The incubation period for *Cryptosporidium spp*. is 2-10 days with an average period of 7 days (17). Cryptosporidiosis was determined to be the cause of diarrheal symptoms as *Cryptosporidium* was common to samples from both patients and the incubation period was most consistent with this pathogen. Neither subject had a history of Cryptosporidiosis prior to this study.

Both subjects reported five days of diarrheal episodes during acute infection with a frequency reaching 8+ episodes during the initial 1-2 and 3 days for subjects A and B, respectively. Subject A lost 4.6lbs (3.3% of total body weight) during acute illness. After resolution of acute symptoms, Subject A experienced several weeks of anorexia concomitant with a further 2.7lbs of weight loss by the end of the study period. Subject B lost 3lbs (2.5% of total body weight) during acute illness which was recovered within three weeks of diarrhea resolution. Both patients were administered oral azithromycin tablets at a dose of 1 mg. Patient A received two doses 1 and 3 days after symptom onset, while patient B received a single dose 1 day after symptom onset. No additional antimicrobial or supportive therapies were administered.

### Cryptosporidium-specific antibodies experience a rapid decay in circulation while avidity continues to increase

Infection with *Cryptosporidium spp*. was verified by detection of antibodies against known antigens Cp17 and Cp23 (**Figure 1A**). Both subjects generated a humoral immune response peaking at the 3-week timepoint in our study and decaying close to baseline by the final 16-week timepoint. Avidity of this antibody response, a marker of B cell maturation, generally increased over the study period (**Figure 1B**). Avidity is the aggregate strength by which a mixture of polyclonal antibodies reacts with antigen epitopes. Subject A displayed higher avidity of all antibody classes (IgM, IgG, IgA) for both antigens (Cp17, Cp23) at the final 16-week timepoint relative to the 3-week measurement. Subject B displayed higher avidity at 16-weeks compared to 3-weeks for all tests except Cp17 IgM and Cp23 IgA. We compared these results to measurements of one- and two-year-old Bangladeshi children sampled 3 weeks after collection of their first *Cryptosporidium* positive stool sample. The American adult subjects were similar in magnitude of antibody generation and antibody avidity compared to Bangladeshi children (**Supp. Figure 1A-B**). Together these data show primary cryptosporidiosis consistently results in circulating antibody against Cp17 and Cp23 antigens; however, levels rapidly decline starting only a few weeks after infection. In contrast, antibody avidity continues to increase over several months without the necessity of a second *Cryptosporidium* exposure.

**Figure 1.**
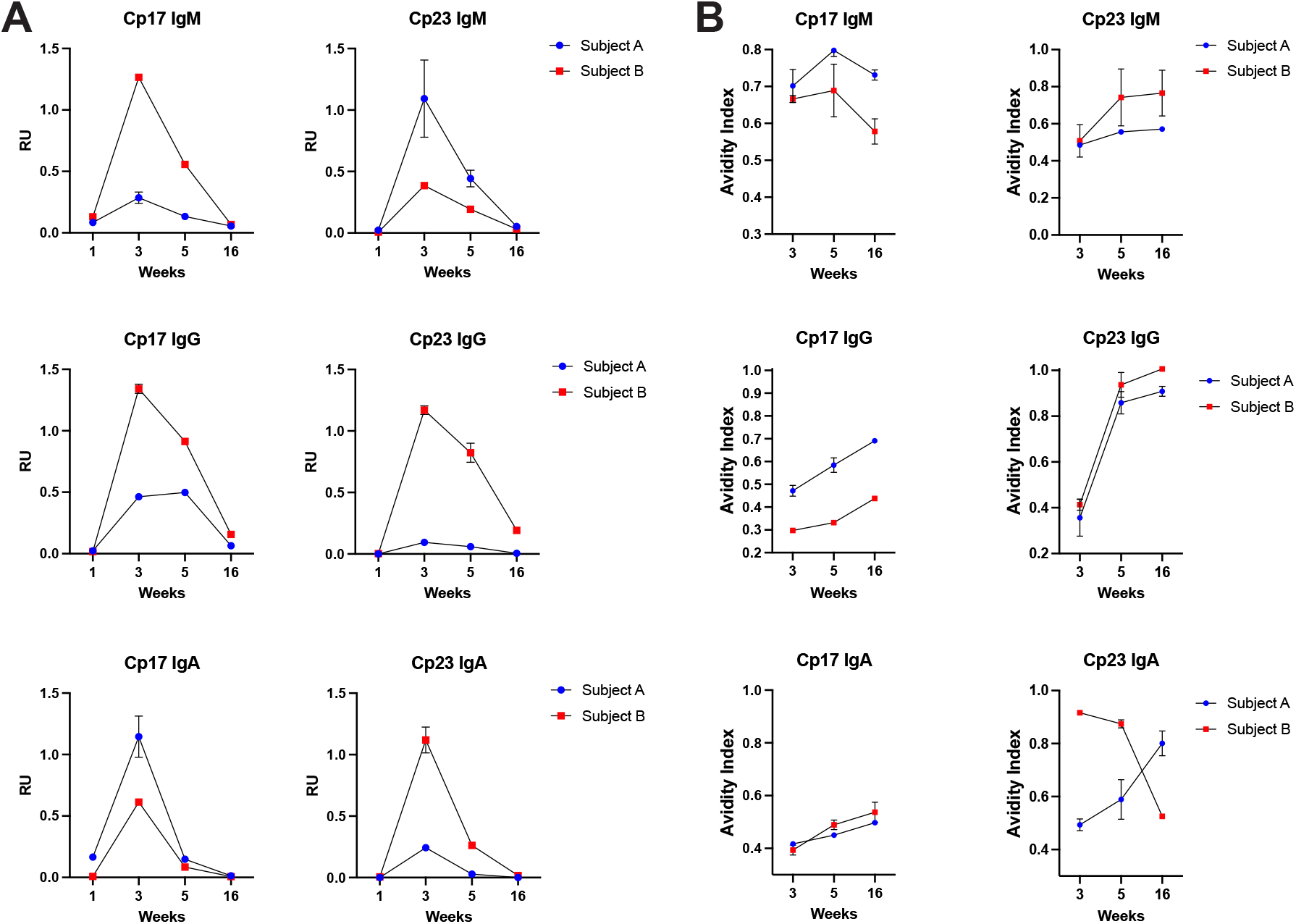
Cryptosporidium-specific antibodies delay rapidly following initial infection while increasing in avidity. **A)** Antibodies specific to *Cryptosporidium spp*. proteins, Cp17 and Cp23, were quantified in subject plasma at weeks 1, 3, 5, and 16 of the study period. **B)** Avidity of antibodies specific to *Cryptosporidium spp*. proteins, Cp17 and Cp23, were calculated at weeks 1, 3, 5, and 16 of the study period. Error bars represent the standard deviation of technical replicates. Error is not shown in instances where the magnitude of the error is smaller than the height of the symbol. RU = relative units. RU calculations are immunoglobulin class-specific and are not directly comparable across sub-classes.

### IgM+ Cryptosporidium-specific circulating memory B cells are dominant post-infection

The quality of antibodies produced following infection is dictated by the differentiation pathway of antigen-specific B cells. Extrafollicular responses rapidly generate plasmablasts and short-lived plasma cells that provide early protection but typically secrete lower-affinity, often IgM-skewed antibodies. By contrast, germinal center responses are sites of somatic hypermutation and affinity-based selection, producing high-affinity, class-switched antibodies (IgG, IgA) and yielding both long-lived plasma cells and memory B cells (18). To investigate the differentiation of Cp17- and Cp23-specific B cells throughout infection, we used fluorescently tagged antigen tetramers to identify our *Cryptosporidium* B cells of interest (**Figure 2A**). Decoy tetramers were used to identify non-specific staining.

**Figure 2.**
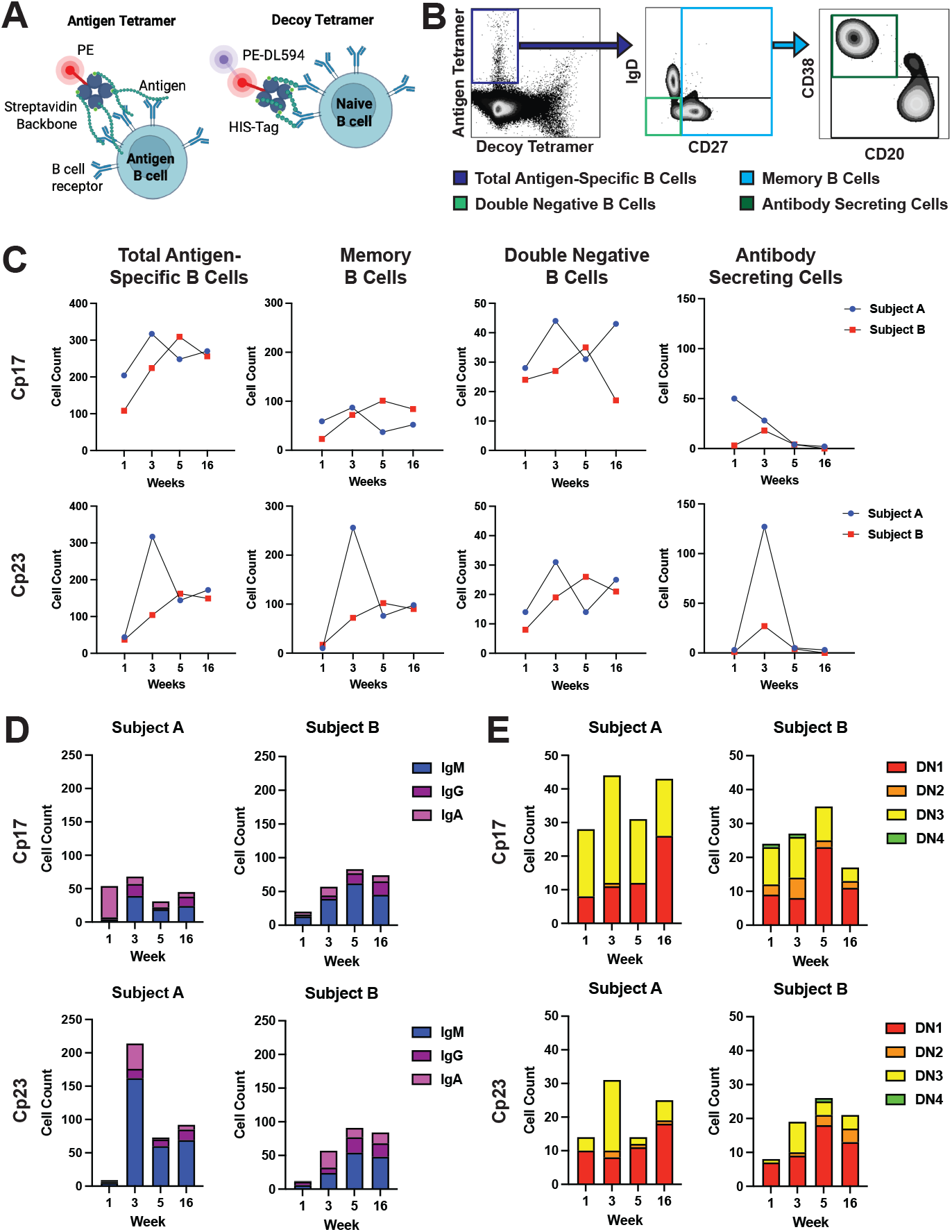
Cryptosporidium-specific memory B cells are predominantly IgM+ post-infection. **A)** Schematic of antigen tetramer (left) and decoy tetramer (right). **B)** Representative gating strategy from PBMCs 3 weeks post infection. **C)** Longitudinal tracking of antigen-specific B cell subsets in subject PBMCs at weeks 1, 3, 5, and 16 of the study period post-infection. Total antigen-specific B cell counts for Cp17- and Cp23 (antigen tetramer+ decoy-), memory B cells (IgD^−^CD27^+^), double negative B cells (DN; IgD^−^CD27^−^), and antibody-secreting cells (CD20^−^CD38^+^) were accounted for after timepoint normalization to total CD45+ cells. **D)** Longitudinal quantification of Cp17- and Cp23-specific memory B cells isotypes (IgM, IgG, IgA) in subjects over time. **E)** Longitudinal quantification of Cp17- and Cp23-specific double-negative B cell subsets (DN1–DN4) in subjects over time. Double negative B cell subsets were determined by CD21 and CD11c expression (DN3, CD21-CD11c-; DN2, CD21-CD11c+; DN1, CD21+ CD11c-; DN4, CD21+ CD11c+).

Cp17- and Cp23-specific B cells (antigen tetramer+ decoy tetramer-) were identified in both subjects (**Figure 2B**). Across the 16-week period post-infection, antigen-specific B cells counts fluctuated, peaking at weeks 3 for Subject A and week 5 for Subject B for both antigens, indicative of an expected early expansion phase of the B cell response and subsequent waning by week 16 (**Figure 2C**). These results did not correlate with changes in the overall B cell population as a percentage of all PBMCs over the course of infection (**Supp. Fig 2**). Analysis of B cell counts by subset indicated differential kinetics. Memory B cells (IgD-CD27+) presented a similar temporal pattern to total antigen-specific B cell counts, peaking around either week 3 or 5 (**Figure 2C**). Double negative (DN) B cells (IgD-CD27-), which have been increasingly linked to chronic inflammation (19), remained relatively stable over time, with minor peaks around week 3 or 5 and a small increase in Subject A or small decrease in Subject B, by week 16 (**Figure 2C**). Antibody secreting cells (CD20-CD38+) were transiently elevated at week 3, consistent with an early plasmablast response following acute infection (**Figure 2C**).

We next conducted an isotype analysis of antigen-specific B cells as a metric of class switching, which could not previously be determined by data presented in **Figure 1A** since relative unit (RU) calculations are immunoglobulin class-specific, and therefore not directly comparable across sub-classes. Antigen-specific isotype analysis of memory B cells unveiled IgM dominated responses, particularly at week 3 in Subject A and in both subjects after week 3. IgG isotypes increased later post-infection, suggesting increased class-switching over time, whereas IgA isotypes were centered around acute infection at week 3 in both subjects and antigens (**Figure 2D**). These patterns were more pronounced for Cp23 than Cp17. Notably, DN B cell subsets 1 and 3 proportionally dominated throughout all timepoints. DN1 B cells peaked at week 16 for Subject A and week 5 for Subject B whereas DN3 B cells peaked at week 3 in all subjects and antigens (**Figure 2E**).

### Subject-to-subject variation and time from infection shape circulating cytokine profiles

We next quantified circulating cytokines at each timepoint using a Luminex 47-plex panel to understand the immune response to cryptosporidiosis. Thirty cytokines were quantifiable at all timepoints for both subjects and were therefore retained in our analysis. Visualization of cytokine profiles by principal component analysis (PCA) showed samples separate by subject on principal component 1 (PC1) and by timepoint on PC2 (**Supp. Figure 3A**). We assessed which cytokines contributed most to PC1 and PC2 to determine responses that characterize variation between subjects and timepoints, respectively.

PC1 was most influenced by drivers of cell growth and wound repair (**Supp. Figure 3B**). Higher levels of wound repair factors in Subject A compared to Subject B may indicate increased intestinal damage and disease severity, which is consistent with the increased weight loss, prolonged anorexia, and higher C-Reactive Protein (CRP) seen in Subject A (**Supp. Figure 3C**). However, inter-subject variation is difficult to assess given the low sample size of two individuals.

The top cytokines influencing PC2 were CXCL9, CXCL10, and IL-27 (**Figure 3A**). This group of cytokines was increased during the acute infection timepoint at Week 1 in both subjects compared to all later timepoints.

**Figure 3.**
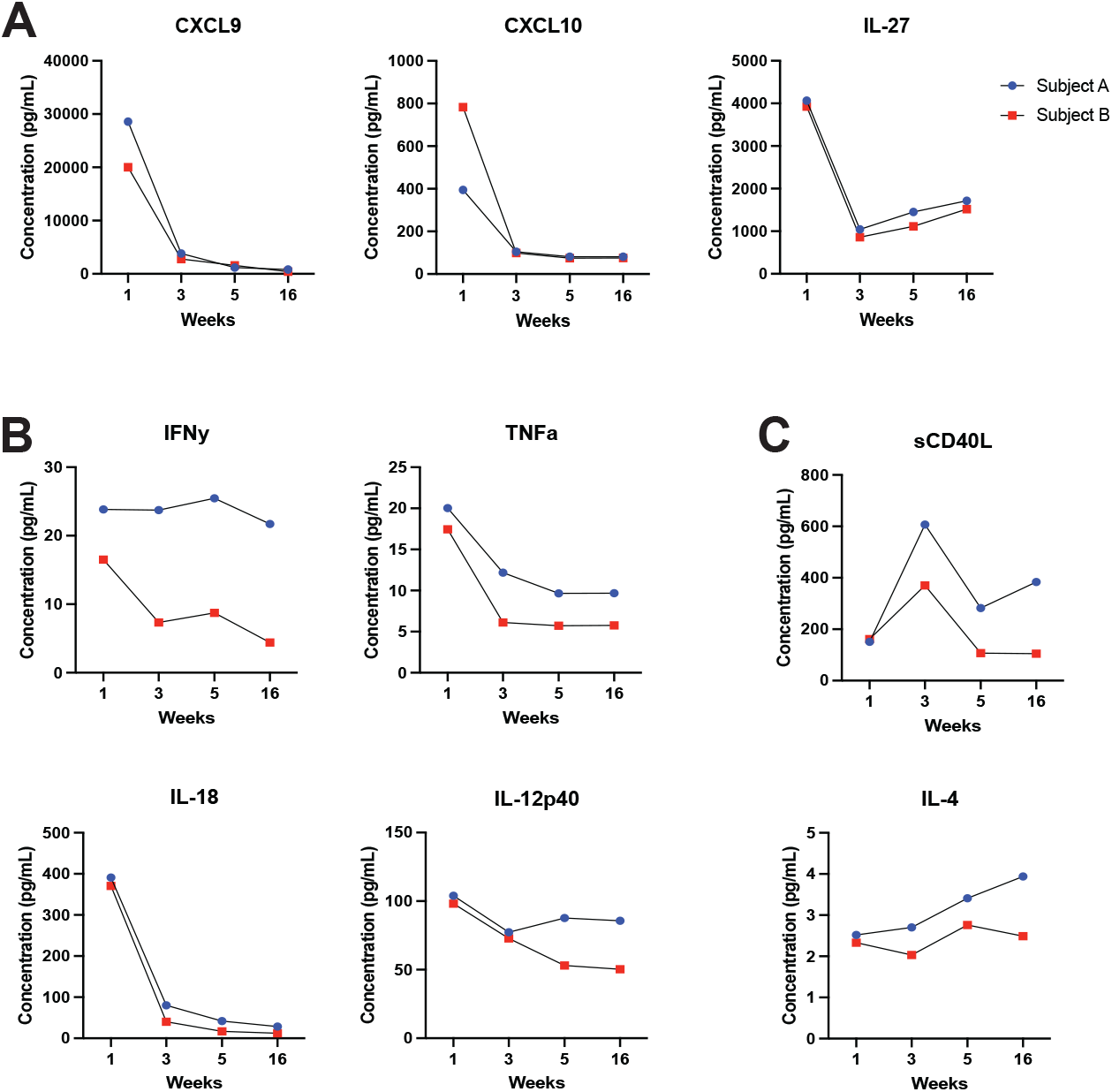
Time from infection identifies Th1 circulating cytokine profiles during acute *Cryptosporidium* infection. **A)** Quantification of cytokines by Luminex bead-based assay over time that contribute most to PC2. Cytokines with previously reported impacts on **B)** innate and **C)** adaptive protection from *Cryptosporidium*.

Notably, CXCL9, CXCL10, and IL-27 are all associated with a type 1 (Th1) immunity signature. Together these cytokine data indicate, that while growth factors may vary between subjects, signaling molecules that drive an increased Th1 response are an important hallmark of the immune response during acute cryptosporidiosis.

### Cytokine profiles of key molecules from murine models of Cryptosporidiosis

We next examined concentrations of cytokines previously implicated in immunity to *Cryptosporidium*. IFNγ is critical for driving defense against the parasite and limiting infection severity in mouse models (6-8, 20). Similarly, inducers of IFNγ, including IL-12 and IL-18, are important for the early response against *Cryptosporidium* (21-23); and in the absence of robust IFNγ production, TNFa has been shown to offer protection in mice *(24)*. In our study, plasma IFNγ peaked early during acute infection in Subject B before waning at subsequent timepoints (**Figure 4A**). In contrast, Subject A retained high IFNγ levels throughout the study period. For TNFa, IL-18, and the IL-12p40 subunit, both subjects measured highest at Week 1 before decreasing at later timepoints (**Figure 3B**).

IL-4 provides a supportive role in parasite clearance by synergizing with IFNγ but additionally modulates the development of acquired immunity by promoting immunoglobulin class switching (25, 26). Binding of CD40 to its cognate ligand also aids in development of acquired immunity, and loss of this interaction in mice leads to inability to clear *Cryptosporidium* (27, 28). In our study, soluble CD40 ligand (sCD40L) experienced a peak in both subjects at the 3-Week timepoint while IL-4 remained constant or slightly increased over the study period (**Figure 3C**).

## Discussion

We quantified the durability and functional quality of circulating antibody production in healthy adults to assess the humoral immune response to primary cryptosporidiosis. In parallel, we assayed cytokine production as a proxy of cell-mediated immunity to gauge how human responses compare to those seen in mouse models of *Cryptosporidium* infection. The major findings from this work are the 1) observation of a rapid decay in circulating *Cryptosporidium*-specific antibody after an initial peak, 2) proportionally IgM dominant memory B cells across all time points, and 3) corroboration of relevance of the Th1 cytokine response seen in mouse models for informing the course of a natural human infection.

### Cryptosporidium-specific antibodies are short-lived in circulation

The longevity of circulating antibody and memory B cells against viral and bacterial antigens can be long-lived. For example, circulating IgG SARS-CoV-2 spike protein titers were largely stable from 3 weeks after Covid-19 symptom onset out to approximately 8 months (29). Circulating IgG and memory specific B cells to vaccinia virus were detectable up to 60 years post-vaccination (30). Similar to these viral pathogens, the bacteria *Yersinia enterocolitica*, a common causative agent of gastroenteritis, resulted in long circulating antibody half-lives for IgM, IgG, and IgA that decayed slowly over 36 months (31). In contrast, humoral immunity against protozoan parasites is generally considered to be short-lived. Antibody responses to *Plasmodium falciparum* infection have been reported to rapidly decline within a few weeks of infection resolution (32, 33). Similarly, circulating antibody against *Cryptosporidium* antigens Cp17 and Cp23 failed to persist at high levels in our subjects. Instead, we observed an increase in antigen-specific immunoglobulin peaking at 3-weeks post infection followed by a rapid decline that was notable just 5-weeks post infection.

While plasma antibody concentrations began to wane a few weeks post infection, antibody avidity continued to increase in the absence of re-exposure. Avidity of antibodies is expected to increase over time after exposure due to do affinity maturation to better target pathogen epitopes. The duration over which avidity increases and reaches a plateau after a single exposure varies depending on the pathogen and individual factors. However, this parameter typically matures gradually over several months following the primary infection, then stabilizes without further increase unless there is subsequent exposure (34-36). Consistent with this our subjects had higher avidity 5-weeks post infection compared to 3-weeks post infection in all cases, and higher avidity still at 16-weeks post infection in most cases. These data indicate that while circulating antibody concentrations decline after primary cryptosporidiosis more rapidly compared to many infectious contexts, avidity maturation in germinal centers remains present.

### Cryptosporidium-specific IgM+ circulating memory B cells are dominant post-infection

In addition to circulating antibody data, our analysis of antigen-specific B cells illustrated the kinetics of B cell responses after primary infection. Cp17- and Cp23-specific B cells expanded early in infection, peaking at weeks 3-5 for each subject before contracting by week 16. This was mirrored in the frequency of memory B cells (IgD-CD27+), whereas double-negative B cells (IgD-CD27-) were comparatively stable, and antibody-secreting cells (CD20-CD38+) were transiently elevated during week 3. These dynamics are typical of an acute response: an early burst of antigen-specific lymphocytes followed by contraction as antigen is cleared (37-39). While this does not reflect the long-lived plasma cells and memory B cells that reside in the tissue that are not captured in peripheral blood, the waning of circulating antigen-specific B cells aligns with the short-lived antibody response we observed and is reminiscent of other apicomplexan infections such as *Plasmodium*, where B cells expand rapidly yet decline without repeated exposure (38).

The phenotypic composition of these B cells was notably skewed toward IgM+ memory B cells. For both Cp17 and Cp23, IgM-expressing cells comprised most antigen-specific memory B cells at all timepoints, whereas IgG+ cells increased only modestly over time and IgA+ cells peaked early. Dominance of IgM+ memory B cells has also been described in other apicomplexan infections; for example, *Plasmodium*-specific IgM+ memory B cells are somatically hypermutated and act as rapid, plastic responders on rechallenge (40-42). Such IgM+ memory B cells can contribute to early parasite control but are often short-lived and may not sustain long-term antibody production, which is consistent with the rapid decline in circulating antibodies we observed. This bias may reflect limited class-switching, potentially due to suboptimal a T follicular helper cell response or cytokine milieu. Our subjects exhibited only modest changes in IL-4 and a transient peak in soluble CD40 ligand, cytokines that drive germinal center reactions and isotype switching (43-45). Therefore, while affinity maturation occurred, the predominance of IgM+ memory suggests that extrafollicular responses may have dominated and germinal center-derived, class-switched memory was limited.

Analysis of DN B cell subsets provides additional insight into the nature of humoral responses. DN3 B cells, which lack CD21 and CD11c and are associated with extrafollicular activation, dominated during acute infection (week 3) but contracted thereafter. In contrast, DN1 cells, which express CD21 and are thought to represent precursors of germinal center-derived memory B cells, were more prominent at week 16 (46). This temporal shift suggests that early responses are driven by extrafollicular pathways, whereas later, a germinal center-like response emerges, possibly contributing to the observed increase in antibody avidity. This limited germinal center response, together with the predominant IgM+ memory B cells, may explain why cryptosporidiosis often fails to induce durable sterilizing immunity and why reinfections are common even in endemic areas.

### Cytokine profiles confirm the importance of the Th1 immune response to Cryptosporidiosis

Studies using controlled human infection models (CHIM) to study Cryptosporidiosis are rare, and the most recent such study utilizing *C. hominis* or *C. parvum* was published nearly two decades ago (47-49). The limited data available from CHIM studies focused on quantifying infectious dose and rate of participant anti-*Cryptosporidium* seroconversion. Epidemiological data on natural human infections have been collected on human cohorts in Brazil, Bangladeshi, Haiti, and other countries (3, 50-53). These cohorts are often comprised of individuals suffering from malnutrition and living in area of high transmission resulting in re-exposure potential over longitudinal sampling. Our study, although low in sample size, offers a unique opportunity to study the development of immunity to *Cryptosporidium* over time in naïve, healthy adults without re-exposure. Our findings largely corroborate results presented from human cohort sampling and transgenic mouse models. Cytokine profiling in our study indicated an acute IFNγ-dependent Th1 response marked by high CXCL9, CXCL10, and IL-27. In other studies of cryptosporidiosis, the concentration of CXCL10 increased in intestinal biopsies of *Cryptosporidium*-infected patients with AIDS compared uninfected patients with AIDS or healthy controls (54). In normal hosts, IFN-γ induced CXCL9 and CXCL10 recruit CXCR3-positive effector cells to control parasite burden (54, 55). While less direct evidence has been presented on the role of IL-27, this cytokine promotes a Th1 immune response through induction of IFNγ. IL-27 is a member of the IL-12 family of cytokines; IL-12 is a key cytokine in defense against *Cryptosporidium* that works in synergy with IL-18 at the intersection of innate and adaptive immunity to drives an IFNγ response (8, 23, 56). Increased CXCL9, CXCL10, IL-27, Il-12, and IL-18 all indicate induction of an IFNγ-driven Th1 response during acute *Cryptosporidium* infection although changes to actual IFNγ concentrations were more subtle.

Th1-associated cytokines and chemokines likely shaped the antigen-specific B cell trajectories seen during this study. IL-12 on its own can directly favor extrafollicular responses and suppress germinal center responses (57). In parallel, IFNγ-induced chemokines CXCL9 and CXCL10 were likely to have guided the migration of CXCR3-expressing B cells and supported plasma-cell diDerentiation in peripheral tissues, possibly favoring rapid IgM output over durable germinal center development (58, 59). By contrast, IL-27, which is important for follicular-helper T (Tfh) cell survival and germinal center maintenance, was scarce after week 1, and only a transient increase in soluble CD40L was observed, coinciding with the peak of antigen-specific B cells, which may have provided a brief window for class switch recombination and affinity maturation (43, 60). Consequently, the cytokine milieu observed may tilt the balance toward rapid, innate-like B cell responses rather than robust germinal center reactions, contributing to the immunological footprint observed in the antigen-specific B cell phenotypes.

Azithromycin treatment may also have contributed to the immune patterns we observed. Beyond its antimicrobial activity, azithromycin’s inhibition of NF-κB and AP-1 signaling in macrophages and dendritic cells would be expected to reduce IL-27 production and limit costimulatory pathways such as CD40/CD86-CD40L, consistent with the transient IL-27 and soluble CD40L responses we observed (61, 62). Additionally, azithromycin directly dampens T cell activation through reduced ICOS, OX40, and mTOR signaling, which can limit Tfh proliferation and germinal center support (63). Azithromycin also suppresses pro-inflammatory cytokines such as IL-12 while enhancing IL-10, creating a regulatory milieu that could restrict the durability of Th1- and Tfh-driven responses (64-66). Such effects would be expected to disrupt sustained Tfh support and germinal center formation, favoring short-lived extrafollicular B cell activation and IgM-skewed antibody output, as seen in our cohort. Thus, while host responses drove immunity to Cryptosporidium, initial azithromycin treatment likely dampened the magnitude and durability of T and B cell responses.

In conclusion, our study provides a rare longitudinal view of the immune response to primary cryptosporidiosis in healthy adults without re-exposure. We demonstrate that circulating *Cryptosporidium*-specific antibody levels rise transiently but wane quickly after infection resolution, while avidity continues to mature over several months, reflecting sustained B cell activity. Concurrently, cytokine profiling revealed a dominant Th1-driven response, marked by IFNγ, CXCL9, CXCL10, IL-27, Il-12, and IL-18, consistent with protective pathways identified in animal models and previous human studies. Detailed phenotyping of antigen-specific B cells showed that Cp17- and Cp23-specific populations expanded early but contracted rapidly, with responses dominated by IgM^+^ memory subsets, suggesting reliance on extrafollicular pathways rather than durable germinal center activity. Supporting this, IL-27 production was short-lived and soluble CD40L signals transient, both pointing to limited Tfh support for sustained class switching. The combination of waning antibody levels, IgM predominance, and brief Tfh-associated signaling highlights constriction in generating long-lived humoral immunity after primary infection. These findings underscore the central role of cell-mediated immunity in controlling infection, while highlighting limitations in the durability of humoral protection after a single exposure. Although based on only two cases, this study emphasizes the value of detailed immunologic profiling in natural human infections and provides a foundation for future work aimed at defining correlates of protection. Such insights are critical for guiding vaccine development and designing strategies to prevent recurrent cryptosporidiosis in vulnerable populations.

## Acknowledgments

We acknowledge members of the Petri, Marie, and Taylor labs at UVA for technical help and thoughtful discussions. We would also like to acknowledge the UVA Flow Cytometry Core Facility cores for technical support. This work was supported by NIH grant AI043596 and a Manning Family Foundation Research Gift, both to W.A.P. A.C.B. was supported a PhRMA Foundation Postdoctoral Fellowship in Translational Medicine.

---

**Supplementary Figure 1.**
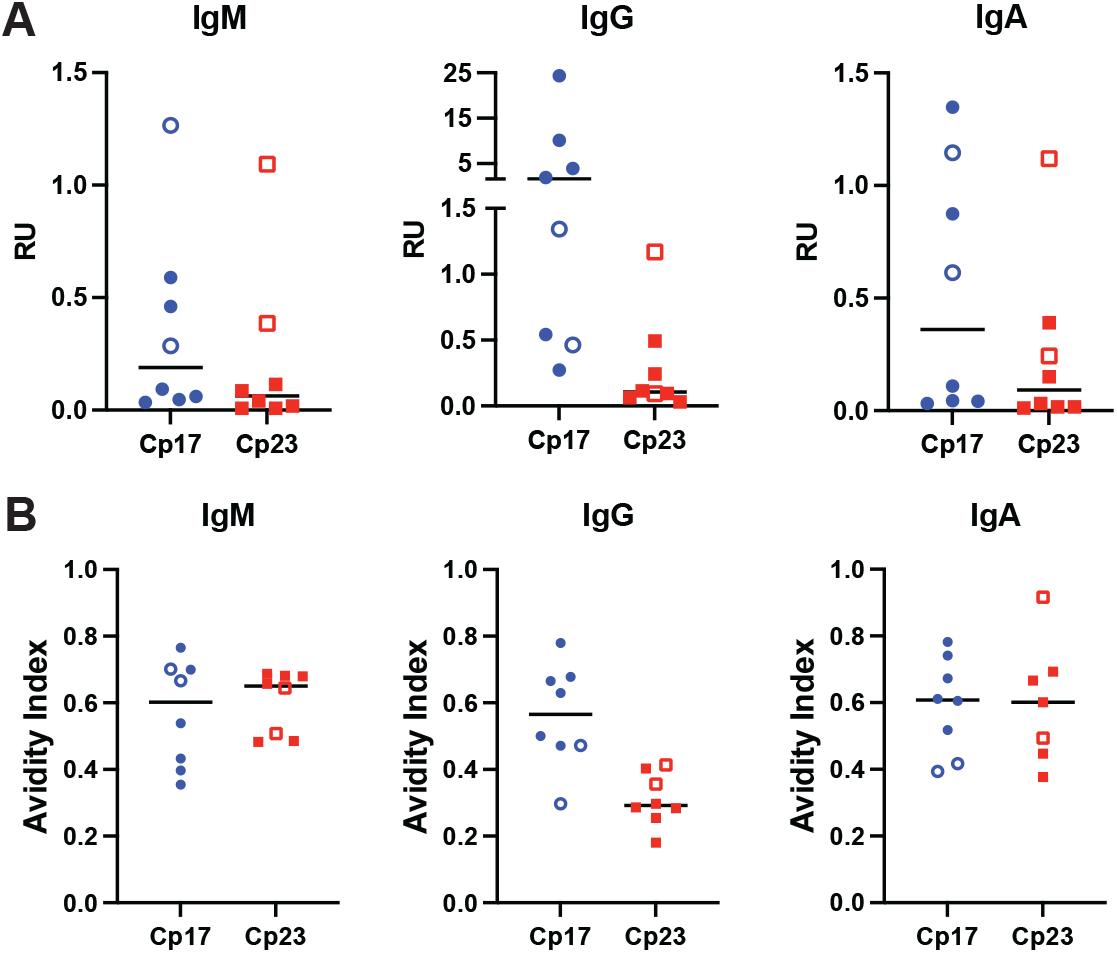
American adults mount a similar antibody response to primary *Cryptosporidium* infection compared to Bangladeshi children. **A)** Antibody quantity and **B)** antibody avidity to *Cryptosporidium spp*. proteins, Cp17 and Cp23, 3 weeks after an individual’s first *Cryptosporidium* infection. Closed symbols indicate Bangladeshi children sampled at one or two years of age. Open symbols indicate American adults. RU = relative units.

**Supplementary Figure 2.**
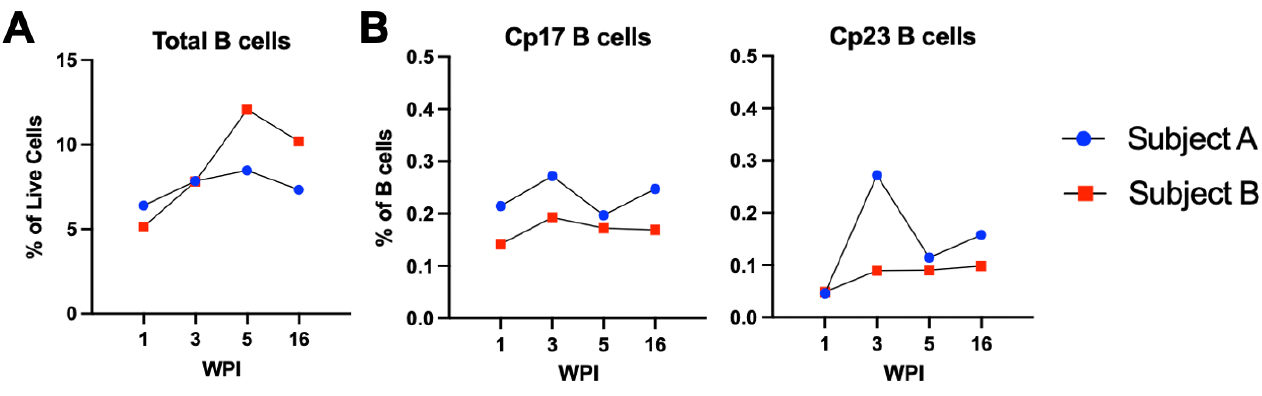
Total measure of B cells increased from start of infection while antigen-specific B cells peak at week 3. **A-B)** Analysis of B cell populations in PBMCs acquired at weeks 1, 3, 5, and 16. **A)** Percentage of total B cell populations (CD3-CD14-CD16-, CD20+ and/or CD19+) out of all live cells. **B)** Percentage of total Cp17 B cells (left) and Cp23 B cells (right) (antigen tetramer+ decoy tetramer-) out of all B cells.

**Supplementary Figure 3.**
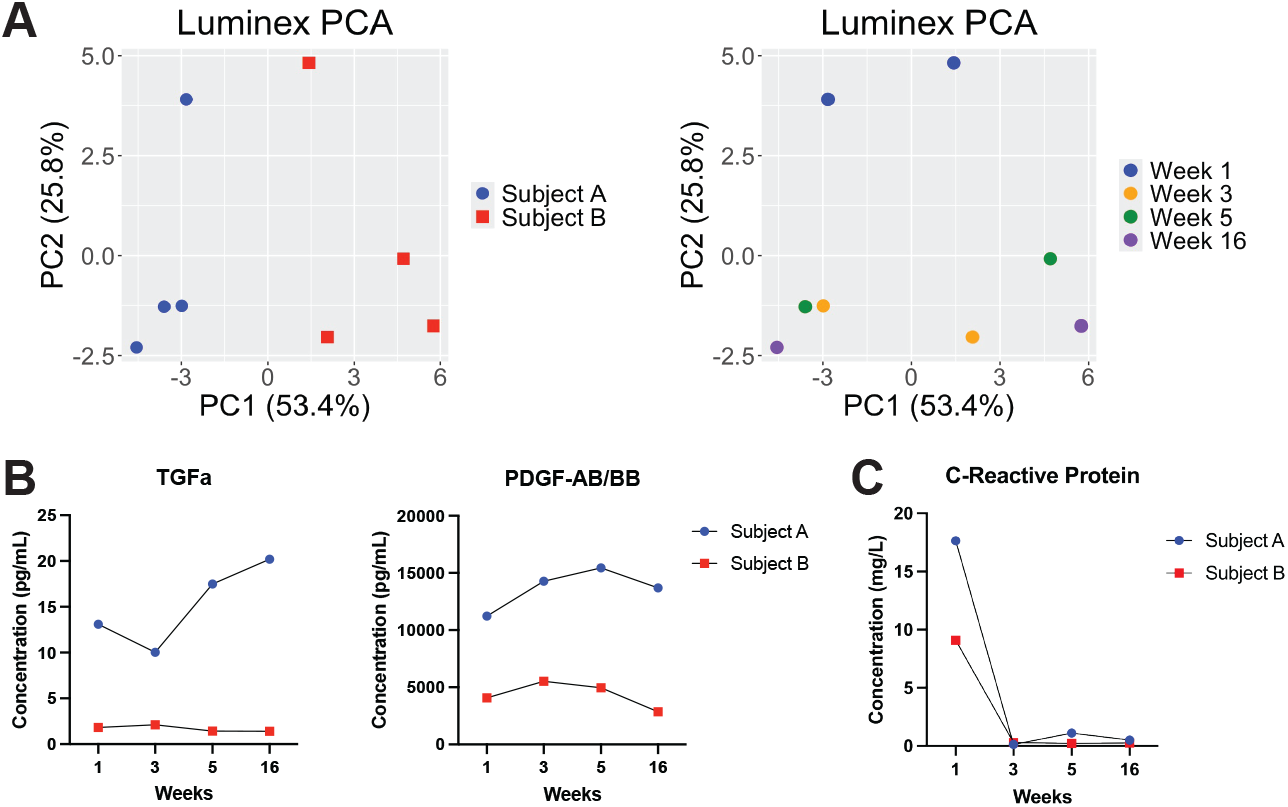
Subject-to-subject variation and time from infection shape circulating cytokine profiles during acute *Cryptosporidium* infection and convalescence. **A)** Visualized principal component analysis (PCA) of plasma Luminex bead-based cytokine quantification color coded by subject (left) and timepoint (right). **B)** Quantification of cytokines by Luminex bead-based assay over time that contribute most to PC1. **C)** Quantification of C-reactive protein generated by enzyme-linked immunosorbent assay (ELISA).

